# Excavating Podocyte-Protective Targets of phloroglucinol-terpene hybrids Utilizing AI Models and Omics Analysis

**DOI:** 10.1101/2025.02.22.639610

**Authors:** Ting Chen, Xiaoyan Liu, Juan Zhu, Xinting Zhang, Jianhui Zhang, Hongping Yu, Ziyan Xu, Dingyan Wang, Yaobin Zhu, Feisheng Zhong, Jiewei Luo

**Affiliations:** School of Medicine, Shanghai University, Shanghai 200444, China; Fujian university of traditional Chinese medicine, Fuzhou 350108, China; Shanghai University of Traditional Chinese Medicine, Shanghai 201203, China; Department of Traditional Chinese Medicine, Shengli Clinical Medical College of Fujian Medical University, Fujian Provincial Hospital, Fuzhou University Affiliated Provincial Hospital, Fuzhou 350001, China; Lingang Laboratory, Shanghai 200031, China; Department of Traditional Chinese Medicine, the First Affiliated Hospital, Fujian Medical University, Fuzhou, China 350005; Fujian Key Laboratory of Drug Target Discovery and Structural and Functional Research, School of Pharmacy, Fujian Medical University, Fuzhou 350122, China.

**Keywords:** PTHs, Podocyte, LN, Llama-Gram model, EGFR

## Abstract

Lupus nephritis (LN) is a multifactorial autoimmune inflammatory disorder, with podocyte dysfunction or structural abnormalities playing a pivotal role in its pathogenesis. *Eucalyptus* leaves, traditionally employed in folk medicine, have demonstrated notable efficacy in managing non-infectious inflammations. Eucarbwenstols D (R14) is a novel phloroglucinol-terpene that has been isolated from *Eucalyptus robusta*. To further refine potential targets, we utilized an AI scoring system and successfully identified the epidermal growth factor receptor (EGFR) as the novel target of R14. This discovery was built upon the foundation of transcriptome and connectivity mapping (CMap) analysis. Molecular docking and molecular dynamics simulations were employed to analyze the interaction between R14 and EGFR, surface plasmon resonance (SPR) and homogeneous time-resolved fluorescence (HTRF) assays were utilized to accurately quantify the binding affinity between R14 and EGFR. Western blot (WB) and immunofluorescence assays were then conducted to explore the downstream signaling pathways initiated by EGFR, specifically the PI3K/AKT/mTOR pathway. These assays demonstrated that R14 modulates the expression of apoptotic proteins Bax and Bcl-2 through this pathway, thereby exerting a protective effect on podocytes from undergoing apoptosis. The modulation of key podocyte proteins, nephrin and podocin, further substantiates this protective effect. Overall, we have applied a novel AI algorithm model that transcends bioinformatics analysis to precise identification of the novel podocyte target of R14. In this study, R14 was identified for the first time as a novel scaffold compound for EGFR inhibition. Moreover, R14 has demonstrated its capacity to inhibit podocyte apoptosis, thereby protecting these critical cells. This discovery establishes a robust theoretical foundation for the use of phloroglucinol-terpene hybrids (PTHs) in the treatment of LN.

## 1. Introduction

Lupus nephritis (LN) is the most severe complication of systemic lupus erythematosus (SLE), affecting approximately 40% of patients and significantly contributing to morbidity and mortality (Ichinose, 2024). Despite improvements in prognosis over recent decades, 5 % – 20 % of patients with LN progress to the end-stage renal disease (ESRD) within 10 years, underscoring the urgent need to identify novel and effective therapeutic targets for LN (Ye et al., 2024). The development of LN is influenced by a variety of factors, including genetic, hormonal, environmental and presence of drugs, can influence the development of LN (Ge et al., 2017). Presently, mycophenolate mofetil, cyclophosphamide, and glucocorticosteroids (GCs) are typically employed as immunosuppressants for the treatment of LN. Nevertheless, the efficacy of these drugs is inconsistent, as 35% of patients may experience relapse after taking immunosuppressive agents. In addition, long-term uses and high-dosing regimen using of immunosuppressants results in toxic side effects, seriously affecting patients’ survival and quality of life. Therefore, there is an urgent need to develop alternative medication that can effectively improve LN and reduce adverse reactions (Yang et al., 2024).

Natural products have long been clinically used and considered as an alternative therapy for the prevention and treatment of various diseases, including LN. In our preliminary work, we found that some compounds isolated from the *Eucalyptus* (*Myrtaceae*) plant *Eucalyptus robusta* have the effect of improving the state of podocytes (Chen et al., 2022). Podocytes are highly specialized and terminally differentiated cells that attach to the outer side of the glomerular basement membrane (GBM) and together with the GBM capillary endothelium, form the glomerular blood filtration barrier. Podocyte play a major role in preventing protein leakage into Bowman’s capsule (Novelli et al., 2018). Podocytes line the urinary surface of the glomerular capillary tuft in the kidneys and play a crucial role in the ultrafiltration of blood. Chronic damaging stimuli, such as infection, metabolic factors and haemodynamic abnormalities, can induce podocyte oxidative stress, leading to podocyte foot-process effacement and loss. Prolonged podocyte injury leads to glomerulosclerosis and the progression of kidney disease (Jin et al., 2018). The majority of LN is glomerulonephritis. Thus, we consider that protecting podocytes is a potential target for treating LN.

In a previous study, Bollée G and colleagues showed that the phosphorylation of epidermal growth factor receptor (EGFR) in podocytes by autocrine heparin-binding EGF (HB-EGF) resulted in the activation of perivascular epithelioid cells (PECs) and development of crescents in a mouse model of anti-GBM disease (Bollée et al., 2011). The phosphorylated EGFR subsequently activates a wide variety of intracellular signaling pathways which are essential for many cellular functions, such as proliferation, migration, differentiation, and apoptosis (Meliambro et al., 2024). These pathways include the mitogen-activated protein kinase (MAPK), janus kinase (JAK) signal transducers and activators of Transcription (STAT), src kinase and phosphatidylinositol-3-kinase (PI3K) pathways.

The PI3K/protein kinase B (AKT) pathway is a major upstream activator of Mammalian target of rapamycin (mTOR) and has been implicated in the propagation of cancer and autoimmunity (Di Cristofano et al., 1999). mTOR is an evolutionarily conserved serine-threonine kinase that regulates cell growth, proliferation, metabolism and survival in response to hormonal and nutrient signals (Weichhart et al., 2015, Fantus et al., 2016). This molecule responds to environmental levels of amino acids, ATP, growth factors, and insulin, performing a critical role in the regulation of cell growth. Under normal conditions, the levels of mTOR in the glomerulus are extremely low, but abnormally high in kidney lesions (Jin et al., 2018). PI3K/AKT/mTOR signaling pathway plays a crucial role in inhibiting apoptosis, promoting cell proliferation, and regulating the expression of inflammatory cytokines. This pathway has been found to be upregulated in lupus B cells and T cells. A recent study, ∼50% of the genes curated as lupus disease genes both from both humans and rodents, could be linked to the mTOR pathway (Stylianou et al., 2011).

Eucarbwenstols D (R14) is a novel PTHs isolated from *E. robusta* in our previously work. R14 showed the promising potential for protecting podocytes (MPC-5 cells) because pre-treatment with pro-inflammatory cytokines TGF-β, IFN-α and IL-6 decreased ROS production and ameliorated the mitochondrial state (Chen et al., 2022). To achieve this goal, we employed transcriptional analysis, Connectivity Map (CMap), and the AI model Llama-Gram to predict the potential target of R14 in podocytes. Llama-Gram represents an improved iteration of the Llama model (Touvron et al., 2023), integrating the Gram Layer to enhance uncertainty estimation (Fan et al., 2024) and effectively mitigate hallucination issues (Farquhar et al., 2024). Leveraging extensive protein-ligand interaction datasets, it is particularly well-suited for tasks involving target prediction.

After predoction, molecular docking, surface plasomn resonanace (SPR) and homogeneous time-resolved fluorescence (HTRF) were followed to validate the binding of R14 and EGFR. Subsequently, western blotting (WB), immunofluorescence and flow cytometry were used to verify the key target pathway. This study aims to investigate the action of PTHs which was isolated from *E. robusta*. A new AI model which named Llama-Gram was utilized to aid the discovery of potential targets of R14. This is a first such use of AI Llama-Gram model. When combined with computer simulation techniques such as transcriptomics to target the EGFR protein, this study further explores for the first time the possibility of PTHs as innovative EGFR scaffolds and offers initial mechanistic insights into how PTHs contribute to podocyte preservation.

## 2. Materials and methods

### 2.1 Preparation and information of R14

Various separation methods (such as silica gel, medium pressure liquid chromatography and preparation-high performance liquid chromatography) were used to isolate in the PE extract of *E. robusta.* Specific steps refer to Chen et al., 2022. The ^1^H NMR and ^13^C NMR of R14 was shown in Fig. S1 and S2.

### 2.2 Cell culture and treatment

The MPC-5 cells were obtained from the Institute of Bio-chemistry and Cell Biology, China Academy of Sciences. Cells were cultured in complete DMEM/F12 (Gibco, USA) medium supplemented with 10% FBS (Biological Industries, South America), at 37 [ in 5% CO_2_ as control group (Con.); MPC-5 cells were treat with immunoglobulin G (IgG; Sigma) for 24 h to establish an *in vitro* model of LN and as model group (IgG); MPC-5 cells in model group treat with R14 in 50 μM (according to Chen et al., 2022).were defined as treated group (IgG+R14).

### 2.3 RNA sequencing

#### 2.3.1 RNA extraction and transcriptome sequencing

MPC-5 cells were treated with 20 µM R14 for 48 h. Total RNA was isolated using Trizol. Each sample was subjected to fragmentation, cDNA synthesis, PCR amplification, and sequenced by the TruSeq Stranded mRNA LT Sample Prep Kit (KP701, Illumina). RNA-seq analysis was completed by H.Wayen Biotech Co., Ltd (Shanghai, China). Differentially expressed mRNAs were selected on the basis of adjusted *p* < 0.05 and |log2fold change|≥0.58. Gene Ontology enrichment and Kyoto Encyclopedia of Genes and Genomes analysis of Differentially expressed mRNAs were achieved using R based on the hypergeometric distribution, respectively.

#### 2.3.2 Bioinformatic analysis

In order to identify internal associations in target sets, we took the intersection of differentially expressed genes (DEGs) into Cytoscape 3.7.2. to construct a Protein-Protein Interaction (PPI) network based on the Search Tool for the Retrieval of Interacting Genes (STRING) databases. Then, modules in protein interaction networks were identified by MCODE algorithm, and the parameters of the MCODE used in this study were as follows: the degree of cut-off, 2; cluster finding, haircut; node score cut-off, 0.2; k-core, 2; and the maximum depth, 100. Furthermore, the CytoNCA plug-in was used to determine network topology parameters, including Betweenness Centrality (BC), Closeness Centrality (CC), and Degree (De), of the PPI modules.

### 2.4 CMap analysis

The gene expression profile of R14 was obtained from our previous dataset (https://www.ncbi.nlm.nih.gov/geo/). The differential expression probes of R14 were selected according to fold-change (FC R 2). The 235 gene expression signatures of R14 were represented by two sets (‘up-’ and ‘down-’ probe sets, saved as .grp files and required as the inputs for CMap), which were made up by the significant up/down regulation probes.

### 2.5 The fold-based AI target prediction methods

To swiftly identify the target of the small-molecule compound R14, the optimized Llama-Gram model was employed for target prediction. The model utilized the ESM-Fold embedding of potential targets along with the graph representation of R14 as inputs. The Llama component of the model analyzed potential intermolecular interactions between the drug and its targets, while the Gram layer provided uncertainty estimation. AI-driven scoring, ranking, and screening were conducted on the 11 potential targets.

### 2.6 Molecular docking of R14 to EGFR

Molecular docking encompasses several key steps: preparing the protein, generating the grid, preparing the ligand, and conducting the molecular docking.

#### Protein Preparation

To prepare the protein, the first step involves downloading the 3D structure of EGFR from the RCSB PDB database ( PDB ID 6LUD). Following this, the protein undergoes processing through a protein preparation module, which is divided into three main stages: Import and Process, Review and Modify, and Refine. The Import and Process stage include correcting bond orders, adding hydrogens, incorporating zero-order bonds to metals, supplementing disulfide bonds, and eliminating water molecules that are more than five angstroms away from the complex structure. In the subsequent Review and Modify stage, molecules that do not contribute to docking are removed, while ensuring the ligand maraviroc is retained. In the Refine stage, the PROPKA program was used to determine the protonation states of protein residues and to optimize the hydrogen bonds that are formed. Finally, the protein structure undergoes energy minimization using the OPLS2005 force field. This process aims to achieve an RMSD of heavy atoms within 0.3 Å, ensuring the structure is optimized for subsequent analyses or simulations.

#### Grid Generation

We use the Receptor Grid Generation module in the Maestro toolkit to generate the protein grid. Through the graphical interface of the Maestro program, we select the small molecule maraviroc in the complex and limit the size of the grid to a 20 Å × 20 Å × 20 Å box centered on the ligand centroid.

#### Ligand Preparation

The R14 compounds undergo processing via the LigPrep module, which performs hydrogenation, energy optimization, and other preparatory steps, resulting in 3D structures ready for virtual screening.

### Induced Fit Docking

The Induced Fit Docking module facilitates flexible docking by importing prepared compounds and protein.

### 2.7 Molecular dynamics simulations

Molecular dynamics analysis was conducted using the Desmond to simulate the complex, incorporating a membrane environment and employing the standard OPLS 2005 force field and the TIP3P water model. Energy minimization and equilibration steps were executed using default settings to achieve a stable initial configuration for the MD simulations. The simulated complex was subjected to a temperature of 298 K and a pressure of 1.01325 bar, which closely experimental conditions. Additionally, appropriate ions were added to mimic the ionic strength of the system.

### 2.8 SPR analysis

The interaction between EGFR protein with R14 was monitored by SPR using a BIAcore T200 (Cytiva) carried out at 25 [ in LMW multi-cycle -cycle mode. Experiments were performed by Wayen Biotechnologies, Inc. The CM5 biosensor chip (Cytiva) was first immobilized with EGFR protein, according to manufacturer’s amine-coupling chemistry protocol (Cytiva). R14 used for this assay were in buffer containing PBS-P (pH 7.4). Various concentrations of R14 was then flowed through the chip and the realtime response was recorded. The concentrations of R14 were 0 nM, 195.3 nM, 390.625 nM, 781.25 nM, 1.56 μM, 3.123 μM, 6.25 μM, 12.5 μM, 25 μM, 50 μM and 100 μM. The equilibrium dissociation constants (binding affinity, KD) for each pair of interaction were calculated using BIAcore T200 evaluation software (Cytiva).

### 2.9 Inhibition of EGFR tyrosine kinase activity HTRF assay

Incubate a series of gradient concentration R14 with a specific concentration of enzyme solution at room temperature for 5 min, add an appropriate amount of enzyme reaction substrate and ATP to start the enzyme reaction process. After 30 min, add a mass of reaction termination solution and detection solution to the enzyme reaction system. Incubate 1 hour, and measure the enzyme activity at the specific R14 concentration on the Flexstation III multi-function measuring instrument. Calculate the inhibitory activity of different concentrations of R14 on enzyme activity, and then fit the inhibitory activity of different concentrations of compounds according to the parameter equation to calculate the value of IC_50_. The EGFR^WT^ kinase and ATP were purchased from Sigma Aldrich (St. Louis, MO, USA). HTRF KinEASE-TK was purchased from Cisbio Bioassays (Codolet, France).

### 2.10 Flow cytometry

MPC-5 cells were seeded at a density of 5 × 10^5^ per well in 6-well plates and treated with R14 for 24 h. Both floating and adherent cells were collected for further analysis. An Annexin V-FITC/PI apoptosis detection kit (BD Biosciences, CA, USA) and propidium iodide (PI) staining (Sigma Aldrich, St. Louis, MO, USA) was used to detect apoptosis, and apoptosis analysis kit (Beyotime, Haimen, China) was used for analysis. Specific operations followed the manufacturer’s instructions. All of them were measured by flow cytometry (Beckman Coulter, USA) following the manufacturer’s instructions.

### 2.11 Immunofluorescence

Differentiated podocytes were fixed with cold methanol at -20° C for 20 min and permeabilized with 0.1% Triton X-100/PBS. After they were washed three times with cold PBS. The cells were blocked with 2% bovine serum albumin (BSA) for 1 h at room temperature and incubated with antibody at 37°C for 2 h or 4°C overnight. Then, the cells were washed three times with PBS and incubated with a secondary antibody conjugated to DyLight 488 or DyLight 594 (Earthox, Millbrae, CA, USA) at 37° C for 1 h, respectively. Nuclei were counterstained with DAPI. The coverslips were mounted onto glass slides, and the images were viewed with an Olympus BX43F fluorescence microscope (OLYMPUS, Japan).

### 2.12 Western blot

Cell homogenates were prepared. Protein samples (30-80 μg) were subjected to 10% sodium dodecyl sulfate-polyacrylamide gel electrophoresis, and transferred onto polyvinyldene fluoride membrane (Bio-Rad Laboratory, Hercules, CA). After incubating in blocking buffer (5% milk in tris-buffered saline containing 0.05% Tween 20) for 1.5 h at room temperature, membranes were incubated with different primary antibodies overnight at 4 °C. Antibodies for p-EGFR/EGFR, p-AKT/AKT, p-PI3K/PI3K, p-mTOR/mTOR, Bax and Bcl-2 were purchased from CST (Danvers, MA, USA) podocin, nephrin were purchased from Abcam (Cambride, MA, USA). Then membranes were washed in TBST and reacted with secondary horseradish peroxidaseconjugated antibody (Santa Cruz, CA; 1:5000) for 1-2 h at room temperature. Antigen-antibody complexes were then visualized using enhanced chemiluminescence reagents (Bio-Rad, Hercules, CA). The density of the immunoreactive bands was analyzed using Image J software (NIH, Bethesda, MD).

### 2.13 Statistical analysis

The data were presented as mean ± SD and analyzed using GraphPad Prism 8.0. One-way ANOVA or two-way ANOVA was used for differences among multiple groups. ** ^(##)^*p* < 0.01, and * ^(#)^*p* < 0.05 were considered to be statistically significant.

## 3. Results

### 3.1 Structural identification of R14

The R14 showed the characteristics of a formyl-phloroglucinol with a eudesmane-type sesquiterpene (Fig. 1a). Amazingly, a rare tetrahydropyran moiety was identificated from the ^1^H NMR (Fig. S1) and HMBC (Fig. S3) spectra. The detailed identification of R14 can be obtained from Chen et al., 2022.

**Fig. 1.**
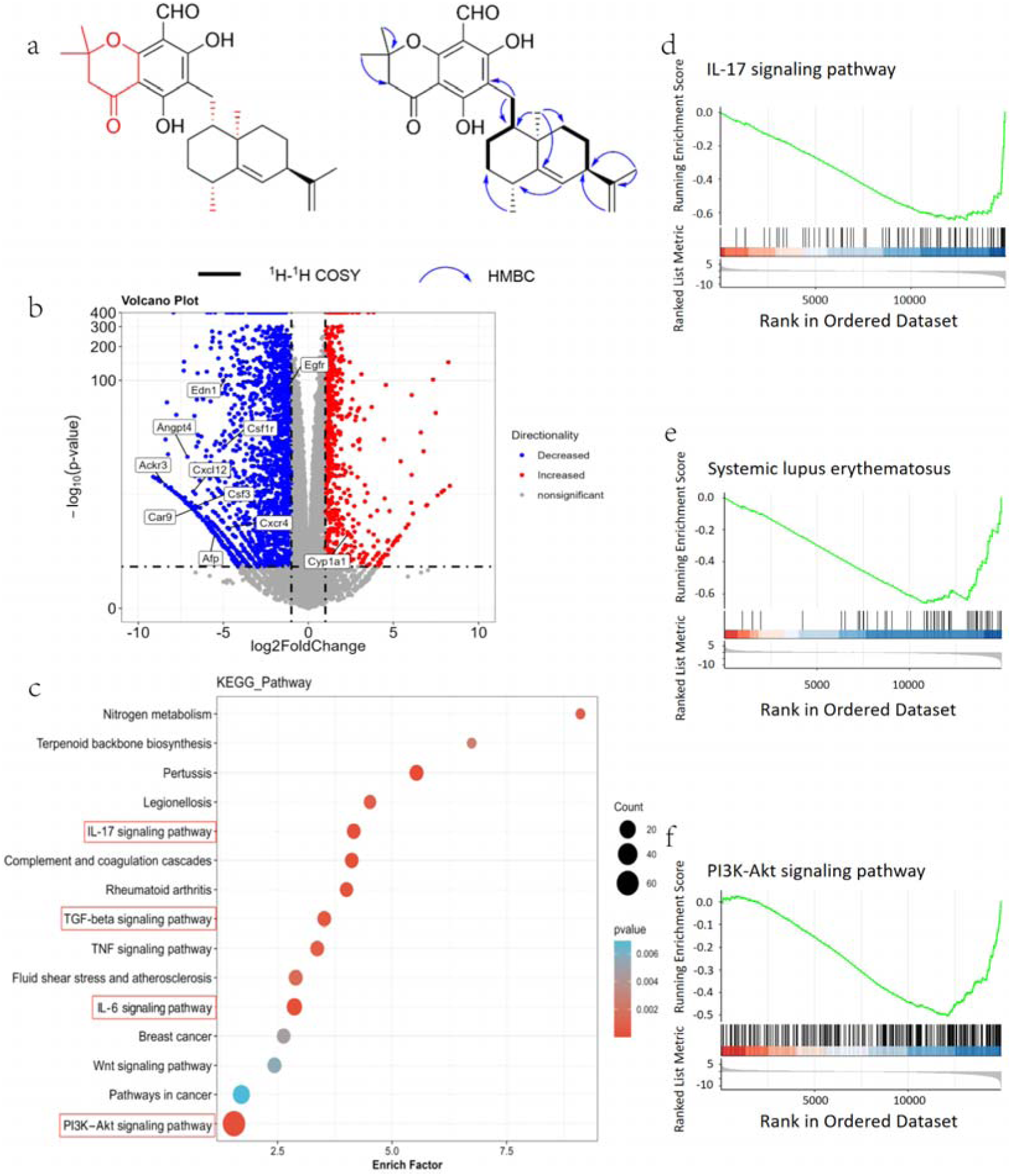
Chemical properties and transcriptome analysis of R14. a. Structure of R14, key ^1^H-^1^H COSY and key HMBC of R14. b. Volcano plot of DEGs in the control and R14 groups; c. KEGG enrichment analyses at the mRNA level; d. Gene Set Enrichment Analysis (GSEA) result pointing to IL-17 signaling pathway in KEGG (Enrichment Score: -0.64607; NES: -1.77014; *p*-value:0.001063); e. GSEA result pointing to systemic lupus erythematosus in KEGG (Enrichment Score: -0.65862; NES: -1.7386; *p*-value:0.001094); f. GSEA result pointing to PI3K-Akt signaling pathway in KEGG (Enrichment Score: -0.50431; NES: -1.46973; *p*-value:0.000999).

### 3.2 Discovery of the potential target of R14 on podocytes

#### 3.2.1 DEGs between R14 and control group

To investigate the potential effects of R14 on MPC-5, we sequenced for mRNA for evidence of variation between conditions of stimulation and non-stimulation. The up-regulated (red) and down-regulated (blue) mRNAs are shown in volcano plot (Fig. 1b). In the decreased part, 10 significantly regulated genes have been identified, among which *Egfr* is prominently listed. The Venn diagram shows the situation of co-expression and specific expression genes, with 260 specific expression genes in the R14 group and 771 specific expression genes in the control group (Fig. S4).

##### DEGs associated with inflammation processes

The KEGG pathway enrichment (Fig. 1c) focuses on the analysis related to inflammatory factors such as IL-17 signaling pathway, TGF β signaling pathway and IL-6 signaling pathway, which are consistent with our previous approach (Chen et al., 2022). Interestingly, PI3K-AKT signaling pathway was enriched with the highest number of genes and statistical significance level. This provided direction for subsequent experimental designs.

In figure 1d-f, IL-17 signaling pathway (Fig. 1d), systemic lupus erythematosus (Fig. 1e) and PI3K-AKT signaling pathway (Fig. 1f) were markedly enriched in the R14 group. These are highly consistent with LN or inflammation related pathways.

The top 10 entries with the highest number of target enrichment ’Counts’ and relatively low *P*-values are shown in GO Terms (Fig. S5). BP included cell surface receptor signaling pathway and anatomical structure morphogenesis. CC were mainly enriched in plasma membrane bounded cell projection, extracellular region and cell projection which indicated that the podocytes may alter their dendritic state or extracellular environment by binding to foreign substances. MF were mainly enriched in small binding, signaling receptor binding and enzyme binding which further supports the information suggested by CC that podocytes may bind to small molecule compounds.

#### 3.2.2 Cmap analysis

DEGs were detected between the R14-treaed and DMSO-treaed groups. Through connectivity mapping (https://clue.io/) visualized the molecular compounds with the most positive correlation with R14 bioactivity in MPC-5 cells were visualized. As shown in Fig. 2a, typhostin-AG-1478 viewed as an EGFR inhibitor have scored 99.73 and its related targets are shown in Fig. 2c. R14 targeted proteins predicted by CMap, the top 11 targets of which are ARF1, ARFGEF1, ARFGEF2, CYTH2, GBF1, SAR1A, EGFR, MAPK14, HSP90AA1, HSP90AA2, and HSP90AB1 (Fig. 2b). However, pinning down a single protein target that is fully responsible for the whole response is not possible. This is because that protein purification and target validation are labor-intensive tasks, and conducting further computational analysis on the predicted targets to narrow down the scope of potential targets becomes essential.

**Fig. 2.**
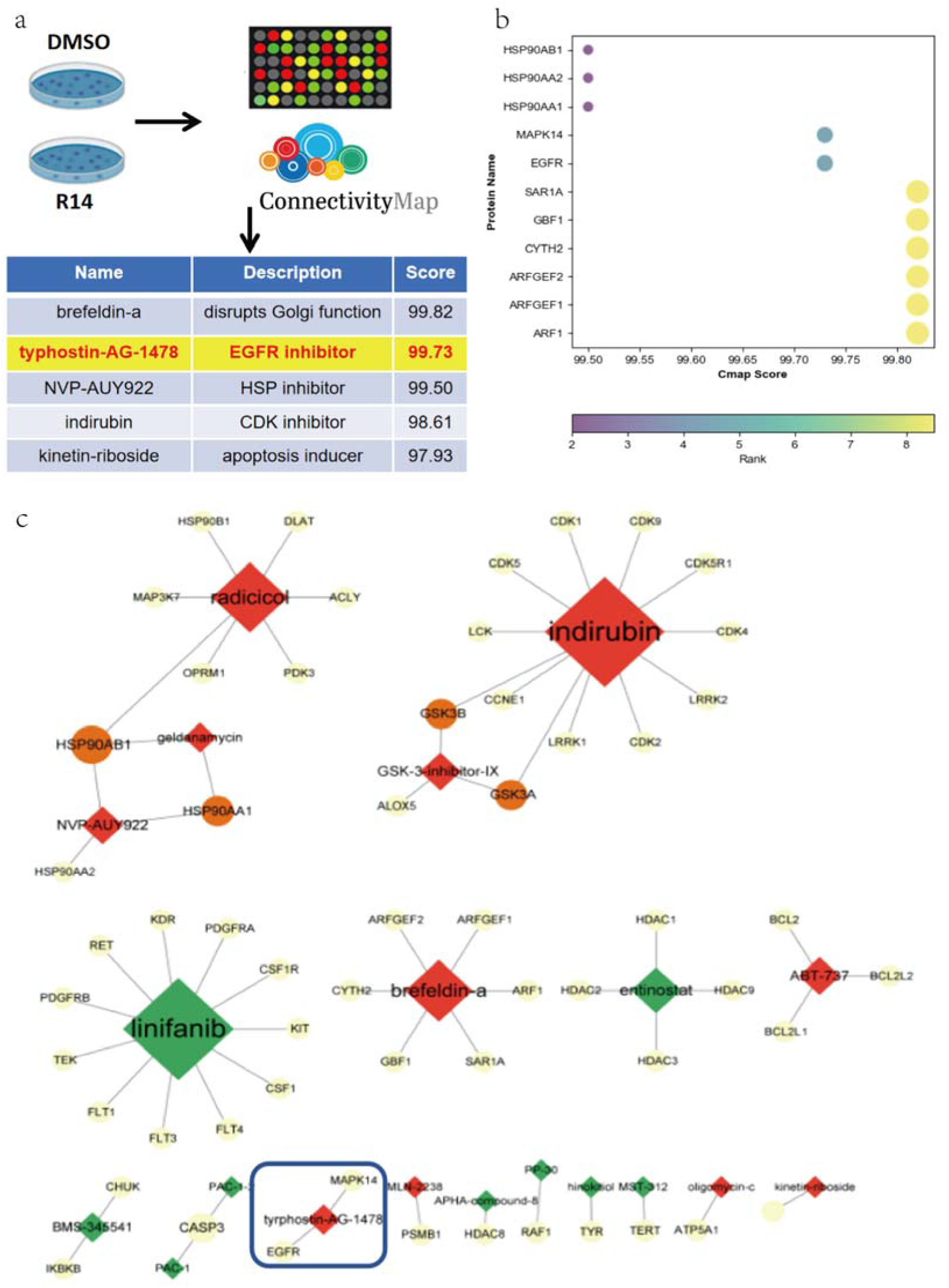
CMap analysis. a. The query genes of R14 were submitted to the CMap database for analysis; b. Top 11 proteins identified through CMap analysis; c. Top 20 compound target relationship network.

### 3.3 Precisely predicting the target of R14 on podocytes using AI model Llama-Gram

To implicitly account for the flexibility of protein conformation, we employed the AI model Llama-Gram for target prediction (Fig. 3a). This model is a primary fold-based, protein conformation-independent, end-to-end deep learning model. The model uses Llama decoding layers for information flow and feature extraction in protein and ligand representation. As shown in Fig. 3b, firstly, the Llama decoding layer adopts RMSNorm normalization to improve the stability and performance of the model. Secondly, the llama decoding layer performs feature interaction and extraction based on the query header (Q) and key header (K). The Llama layer also uses grouped query attention (GQA) mechanism to replace the traditional multi head attention mechanism (Fig. 3c). By sharing K and V pairs, it not only significantly reduces the total number of parameters in the model, but also significantly improves the processing efficiency. The Llama model often suffers from hallucination issues, frequently providing unreasonable estimates. To address this problem, we focus on uncertainty estimation and propose an improved model called the Llama-Gram model to reduce hallucination. Finally, we constructed an in-house protein-ligand dataset comprising 2,577,801 samples to train the Llama-Gram model. Since it is specifically trained for target prediction tasks, it can be effectively applied to target prediction scenarios.

**Fig. 3.**
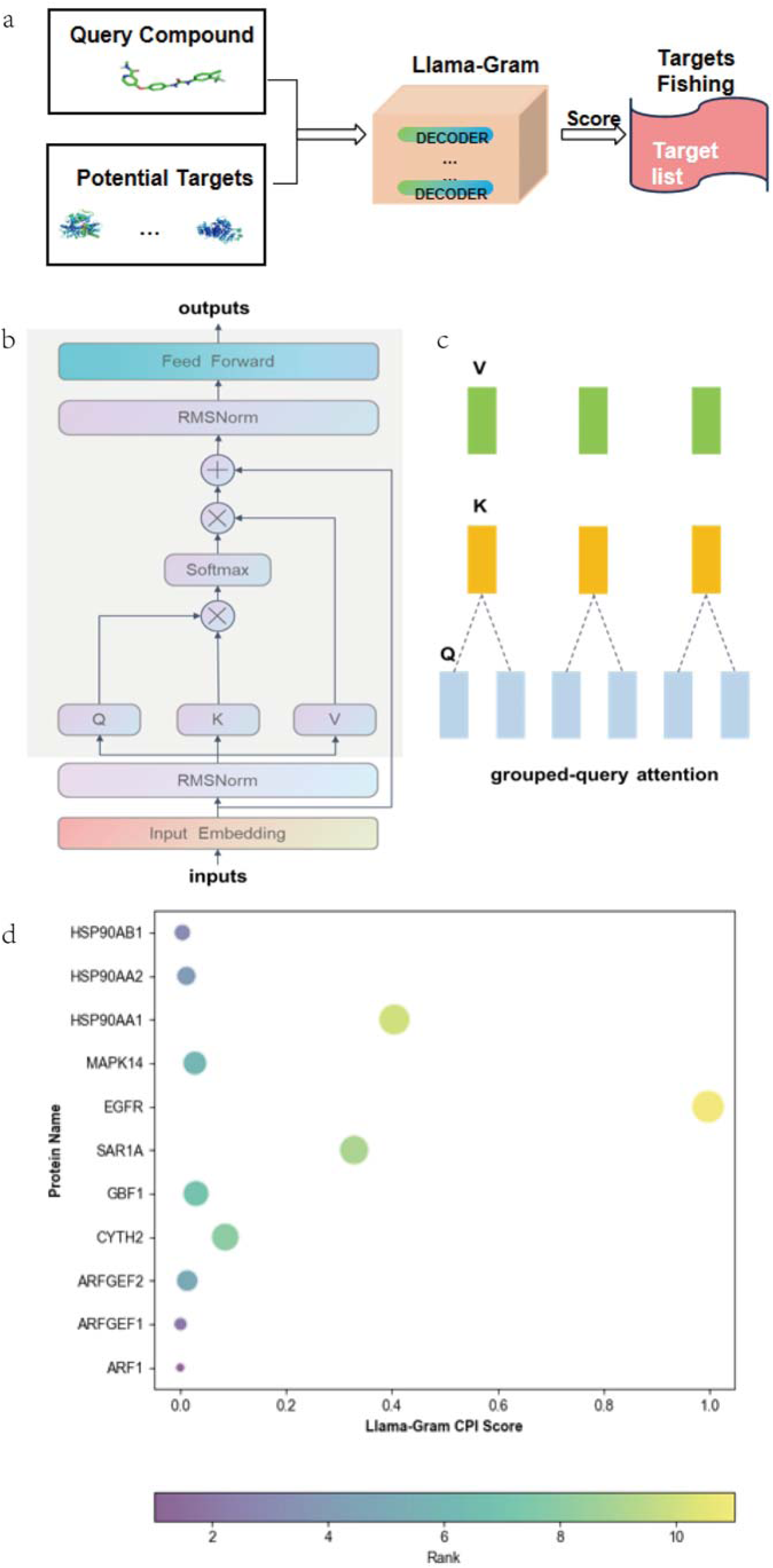
AI model. a. Pipeline of the target inference using the Lllama-Gram model; b. The architecture of the decoder layer; c. Schematic diagram of graph query attention mechanism; d. The bubble chart generated by ranking proteins based on Llama-Gram CPI scores.

The optimized Llama-Gram model was used to evaluate the potential interaction between R14 and 11 potential protein targets. These 11 potential protein targets are ranked in descending order of Llama Gram score (Fig. 3d). Remarkably, the AI model effectively distinguished between the 11 targets from each other that CMap struggled to differentiate. Among these, the Llama-Gram model identified EGFR as the most likely potential target for R14, as it received the highest score. Following upon this important revelation, the wet lab experiments below were conducted to the prediction.

### 3.4 Wet experimental validation and subsequent molecular modeling to establish EGFR as a target of R14

#### 3.4.1 SPR analysis

To determine whether R14 directly binds to EGFR, we examined the direct binding of R14 to purified EGFR *in vitro* using surface plasmon resonance technology. The SPR analysis revealed that the binding of R14 with EGFR exhibited a fast-on and fast-off kinetic pattern (Fig. S8). This finding was not well described with a 1:1 binding model. Therefore, we opted to derive equilibrium dissociation constants using steady state fitting with a K_D_ of 6.0 μM (Fig. 4f).

**Fig. 4.**
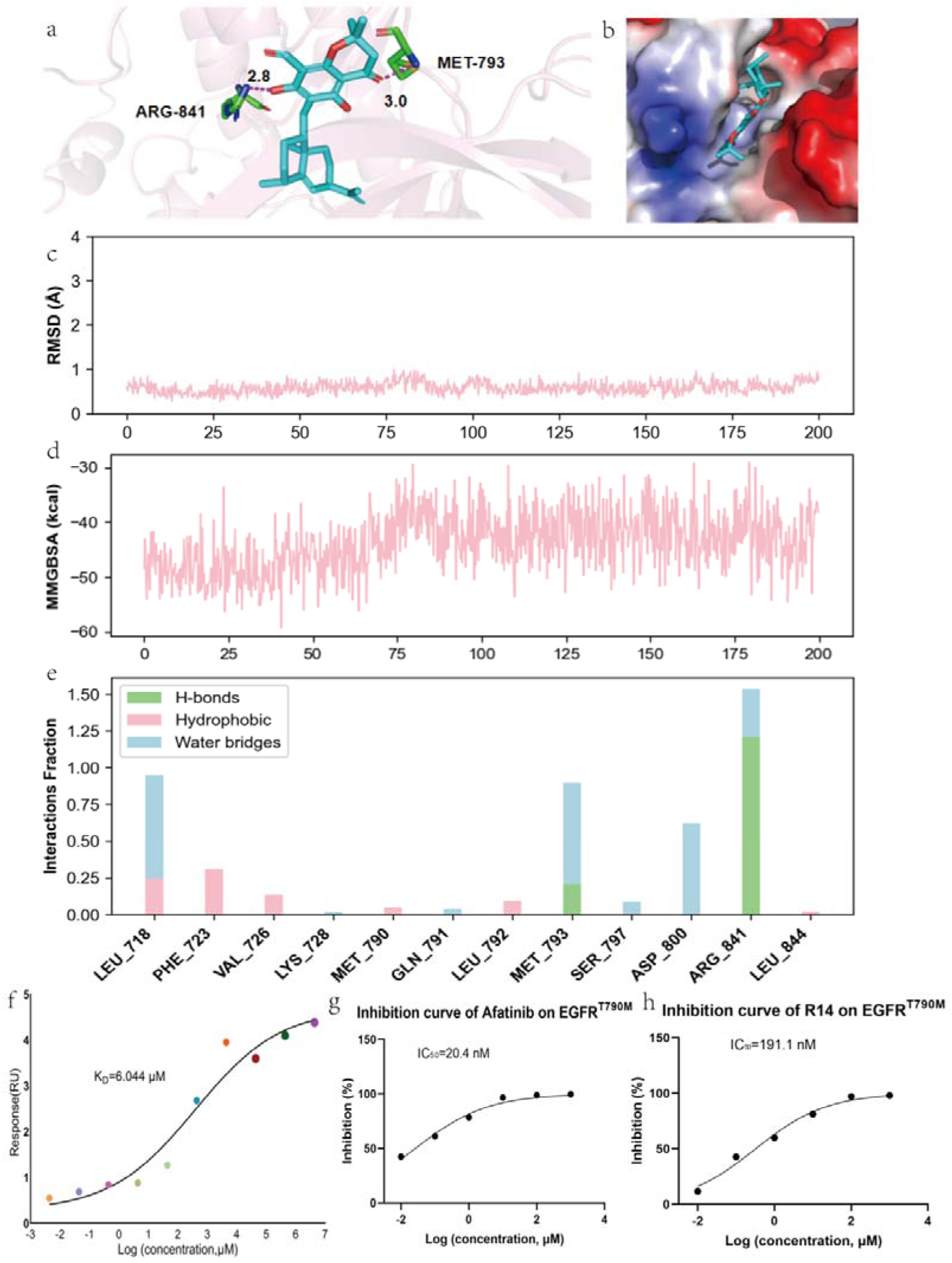
Molecular interaction. a and b. Molecular docking of R14 to EGFR; c. The fluctuation of RMSD over time; d. The fluctuation of MMGBSA over time; e. The interaction fraction of R14 binding sites residues; f. SPR data fitted using a steady state model; g. Inhibition curve of afatinib on EGFR^WT^; h. Inhibition curve of R14 on EGFR^WT^.

#### 3.4.2 HTRF assay

Based on the computational analysis result and the analysis result SPR, we further evaluated the potential of R14 to inhibit the EGFR enzyme activity was evaluated with an *in vitro* HTRF technology. The R14 was identified as an EGFR inhibitor with IC_50_ of 191.1 nM (Fig 4h), even though the effect was weaker than a known EGFR positive inhibitor afatinib (Fig 4g).

#### 3.4.3 Molecular docking and Molecular Dynamics (MD)

Based on the docking pose predicted by Induced Fit Docking, we further conducted 200 ns of MD simulations to elucidate the binding mechanism of R14 in the ATP binding pocket of EGFR (Fig. 4c, d). The hydrophobic functional group of R14 fitted well into a hydrophobic subpocket by PHE-795, PRO-794, MET-793, LEU-792, MET-790 and LEU-788. The carbonyl group on the benzene ring formed a possible strong hydrogen bond with MET-793 (Fig. 4a, b). Importantly, the phenolic hydroxyl group was deprotonated and formed a possible strong hydrogen bond with ARG-841. Furthermore, The RMSD of the ligand remained less than 2, indicating that the conformation of the ligand did not undergo significant changes during the MD simulation process. The MMGBSA method calculated the binding free energy of R14 and EGFR as -43.77 kcal/mol, indicating a strong interaction. The hydrogen bond formed between R14 and ARG-841 remained exceptionally stable during the molecular dynamics process, securely anchoring R14 within the pocket.

Overall, SPR analysis, enzyme activity experiments, molecular docking, and molecular dynamics studies have shown that R14 can form a stable complex with the ATP binding pocket which is consistent with our CMap results.

### 3.5 Regulation of R14 on EGFR downstream signaling pathway

#### 3.5.1 The PI3K/AKT/mTOR signaling pathway

The preceding series of experiments verified the prediction that R14’s podocyte target is EGFR. Our investigation then delved into the downstream signaling pathways initiated by EGFR. The PI3K/AKT/mTOR pathway was scrutinized using WB (Fig. 5a-e) and immunofluorescence (Fig. 6) assays.

**Fig. 5.**
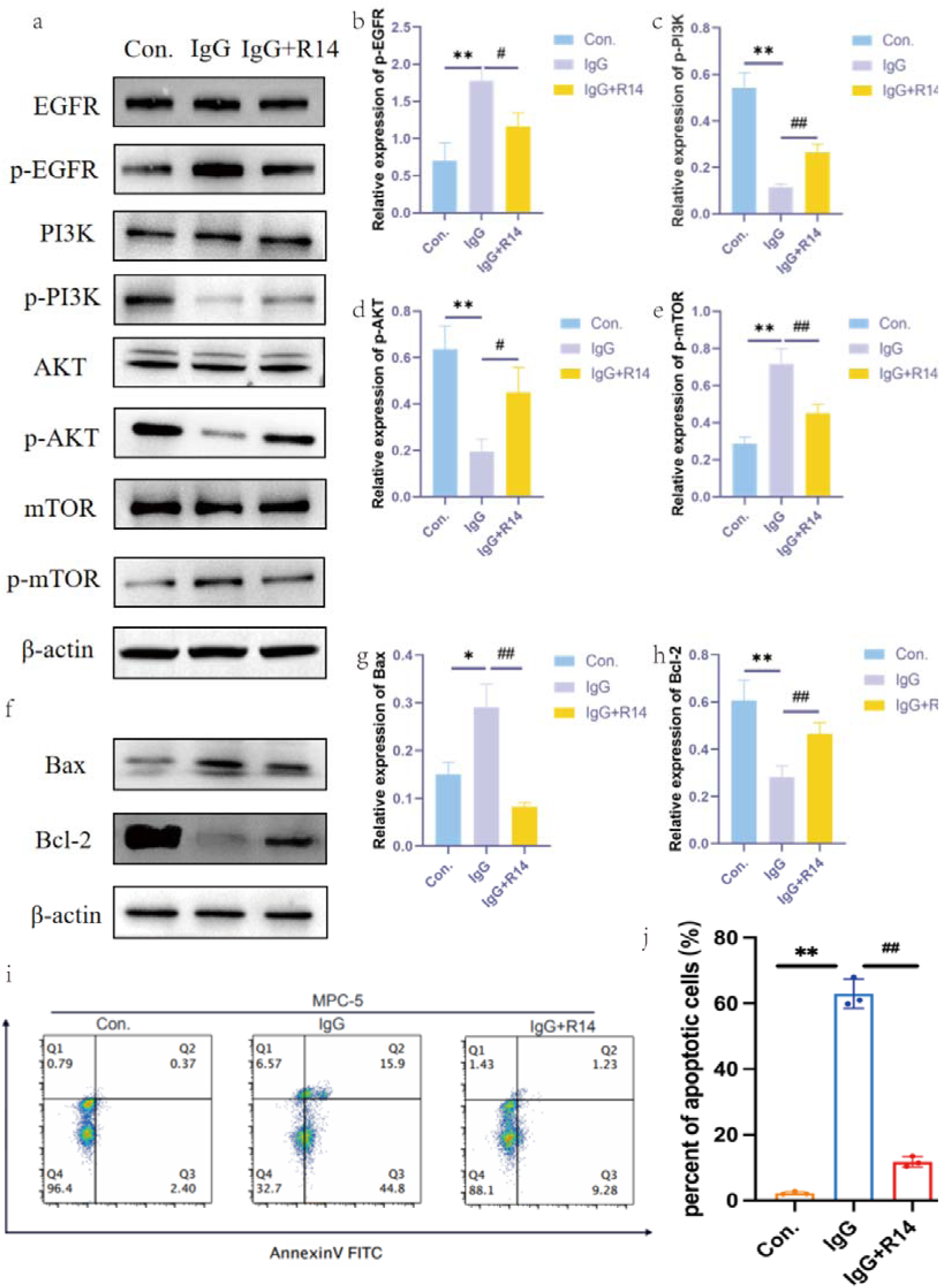
Regulation of R14 on EGFR downstream signaling pathway. a. EGFR/p-EGFR, PI3K/p-PI3K, AKT/p-AKT, mTOR/p-mTOR; b. Statistical analysis of EGFR/p-EGFR. Data were expressed as mean±SD, n=3. \*\**p*<0.01 control vs IgG, #*p*<0.05 IgG vs IgG+R14; c. Statistical analysis of PI3K/p-PI3K. Data were expressed as mean ± SD, n=3. \*\**p*<0.01 control vs IgG, ##*p*<0.01 IgG vs IgG+R14; d. Statistical analysis of AKT/p-AKT. Data were expressed as mean ± SD, n=3. \*\**p*<0.01 control vs IgG, #*p*<0.05 IgG vs IgG+R14; e. Statistical analysis of mTOR/p-mTOR. Data were expressed as mean±SD, n=3. \*\**p*<0.01 control vs IgG, ##*p*<0.01 IgG vs IgG+R14. f. Western blot analysis of Bax and Bcl-2; g. Statistical analysis of Bax. Data were expressed as mean±SD, n=3. \**p*<0.01 control vs IgG, ##*p*<0.05 IgG vs IgG+R14; h. Statistical analysis of Bcl-2. Data were expressed as mean±SD, n=3. \**p*<0.01 control vs IgG, ##*p*<0.05 IgG vs IgG+R14; i and j. The cell-cycle distribution and percentage of apoptotic cells were analyzed by flow cytometry (n=3; \*\**p*<0.01 control vs IgG, ##*p*<0.05 IgG vs IgG+R14).

**Fig. 6.**
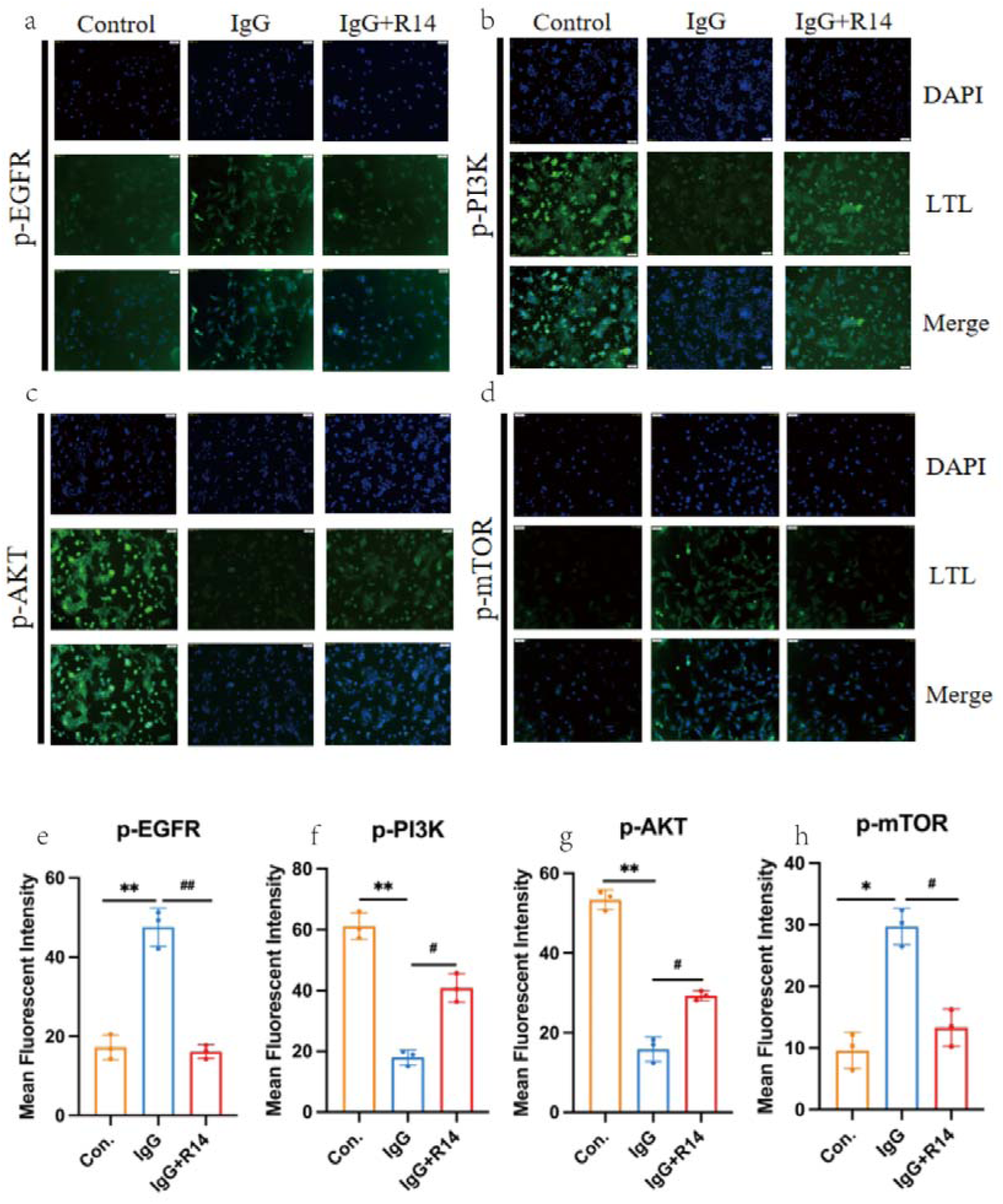
Immunofluorescence analysis for expression levels of p-EGFR, p-PI3K, p-AKT and p-mTOR. a. Images of DAPI/LTL staining double-staining assay of p-EGFR (scale bar: 50 μm); b. Images of DAPI/LTL staining double-staining assay of p-PI3K (scale bar: 50 μm); c. Images of DAPI/LTL staining double-staining assay of p-AKT (scale bar: 50 μm); d. Images of DAPI/LTL staining double-staining assay of p-mTOR (scale bar: 50 μm); e. Bar graph demonstrating the effect of 20 µM, p-EGFR in the presence (n=7) or absence (n<3) of podocyte. **/##*p*< 0.01 and */#*p*< 0.05, paired t test; f. Bar graph demonstrating the effect of 20 µM, p-PI3K in the presence (n=7) or absence (n<3) of podocyte. **/##*p*< 0.01 and */#*p*< 0.05, paired t test; g. Bar graph demonstrating the effect of 20 µM, p-AKT in the presence (n=7) or absence (n<3) of podocyte. **/##*p*<0.01 and */#*p*<0.05, paired t test; h. Bar graph demonstrating the effect of 20 µM, p-mTOR in the presence (n=7) or absence (n<3) of podocyte. **/##*p*<0.01 and */#*p*<0.05, paired t test.

As depicted in Fig. 5a, the WB analysis revealed that the R14 treatment group notably suppressed the IgG-induced upregulation of EGFR phosphorylation and mTOR phosphorylation. Concurrently, R14 significantly enhanced the protein expression levels of PI3K phosphorylation and AKT phosphorylation, which were otherwise diminished by IgG. The immunofluorescence assays provided further validation of the WB findings, as illustrated in Fig. 6.

The PI3K/AKT/mTOR pathway we selected in this experiment is a common downstream pathway of EGFR; At the same time, it worked consistent with our transcriptome prediction analysis (Fig 1c and 1f).

#### 3.5.2 R14 reversed podocyte apoptosis through inhibiting the EGFR pathway

The PI3K/AKT/mTOR signaling pathway is known to modulate the expression levels of the Bcl-2 and Bax proteins, which play pivotal roles in regulating cell survival and apoptosis. WB experiments substantiated this relationship, showing that R14 treatment activated the downstream anti-apoptotic protein Bcl-2 and concurrently suppressed the pro-apoptotic protein Bax, as evidenced in Fig. 5 f-h.

Podocyte apoptosis was assessed using Annexin V staining. The findings indicated that R14 effectively counteracted IgG-induced podocyte apoptosis (Fig. 5i and j). When apoptosis was inhibited, potocytes remained viable. This protective effect was likely attributed to the observed alterations in the expression of Bcl-2 and Bax.

### 3.6 R14 inhibited IgG-induced podocyte injury

As previously discussed, the preservation of podocyte integrity represents a promising therapeutic approach for the management of LN. A hallmark of podocyte dysfunction in LN patients is the diminished expression of nephrin and podocin proteins, as referenced in literature (Maeda et al, 2018, Tian et al., 2020). We quantified the expression levels of these proteins using WB analysis (Fig. 7 a-c) and immunofluorescence (Fig. 7 d-g).

**Fig. 7.**
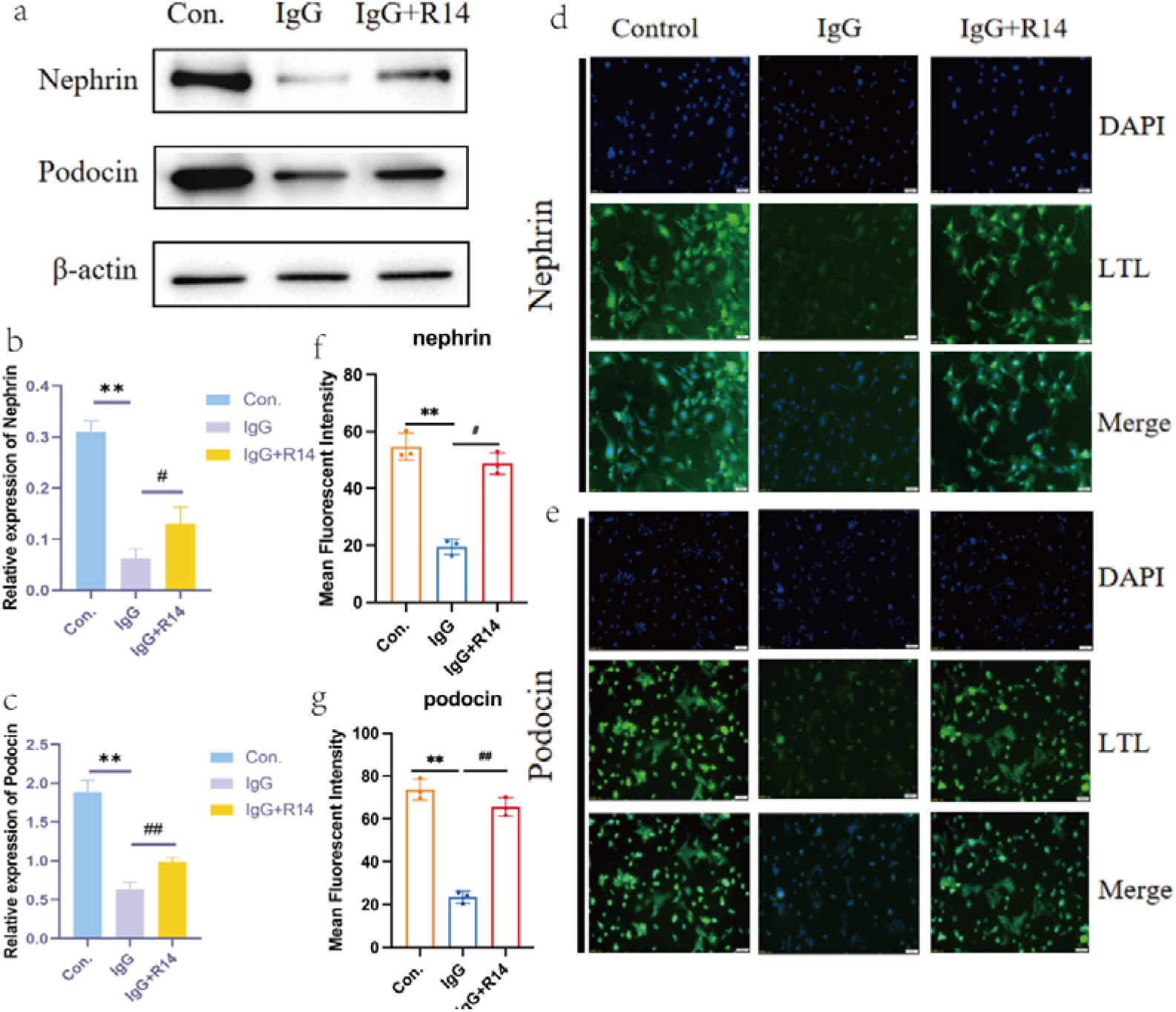
The impact of R14 on key proteins in podocytes. a. Western blot analysis of nephrin and podocin; b. Statistical analysis of nephrin. Data were expressed as mean±SD, n=3. \*\**p*<0.01 control vs IgG, #*p*<0.05 IgG vs IgG+R14; c. Statistical analysis of podocin. Data were expressed as mean±SD, n=3. \*\**p*<0.01 control vs IgG, ##*p*<0.05 IgG vs IgG+R14; d. Images of DAPI/LTL staining double-staining assay of nephrin (scale bar: 50 μm); e. Images of DAPI/LTL staining double-staining assay of podocin (scale bar: 50 μm); f. Bar graph demonstrating the effect of 20 µM, nephrin in the presence (n=7) or absence (n<3) of podocyte. **/##*p*<0.01 and */#*p*<0.05, paired t test. g. Bar graph demonstrating the effect of 20 µM, podocin in the presence (n=7) or absence (n<3) of podocyte. **/##*p*<0.01 and */#*p*<0.05, paired t test.

The data presented in these figures revealed that R14 effectively elevated the reduced expression of podocin and nephrin in IgG-induced podocytes, displaying the ability of R14 in mitigating IgG-induced podocyte injury.

## 4. Discussion

SLE is a chronic autoimmune disease with a diverse course that was difficult to standardize for patient treatment. LN is one of the most common and serious complications of SLE. Currently, there is no complete cure or highly effective medication. GCs can quickly alleviate disease progression and serve as the basic medication for a long term LN treatment, but the administration of higher GCs doses has been associated with organ damage. With the emergence of targeted therapies and the use of small molecular compounds, the FDA has approved Belimumab (Liébana et al., 2024) and Anifrolumab (Jayne et al., 2023) for the treatment of LN; Although monoclonal antibodies have high specificity and can recognize and bind to specific antigenic epitopes; It is precisely because of this feature that they often make it difficult to consider complications. Antimalarials, another clinical medication for LN, can reduce the risk of renal flare, ESRD and death (Alarcón et al., 2019, Pakchotanon et al., 2018); However, the scarcity of data on pharmacokinetics and posology do not allow for indication of recommended dosage, especially in pregnant women.

On one hand, the pathogenesis of LN often encompasses intricate mechanisms, but on the other, natural products (NPs) can act on multiple targets and pathways with complex and viariable mechanisms of action (Chen et al., 2024). *Eucalyptus* have shown treament potentials for various inflammatory conditions (Chen et al., 2022). The leaves of *Eucalyptus* are often used as folk medicine, which has significant curative effect on arthritis, asthma, diabetes etc (Ji et al., 2018). Even though the active compound(s) was unknown to the users and practitioners in ancient times, it were exactly these that presumably exerted the anti-inflammatory effects. Our previous work elucidated that PTHs represented by R14 were active ingredients in this plant material. Another study has indicated that *Eucalyptus* gene remedies may contribute to the prevention and treatment of rheumatoid arthritis by acting on multiple pathways via a variety of active components, especially eucalyptol (Iqbal et al., 2024). Activated B cells, as a type of antigen-presenting cell, promotes the activation of pathogenic T cells to secrete proinflammatory cytokines such as IL-6 and TNF-α, and facilitate the recruitment of macrophages and dendritic cells into the glomerulus and the tubulointerstitium (Liu et al., 2023).

Pyrimidine derivatives substituted with phenylamine, indole, pyrrole, piperazine, pyrazole, thiophene, pyridine and quinazoline derivatives substituted with phenylamine, pyrimidine, morpholine, pyrrole, dioxane, acrylamide, indole, pyridine, furan, pyrimidine, pyrazole etc. are privileged heterocyclic rings shown promising inhibitory activity against EGFR and TKIs (Makhija et al., 2024). Acyl phloroglucinols constitute the largest group of natural phloroglucinols from the *Eucalyptus* genus. The phloroglucinol-terpene adducts are the most common acyl pholoroglucinols reported in the *Eucalyptus* genus. The phloroglucinol-terpene adducts comprise hybrid natural products with high structural diversity that incorporated mono- or sesqui-terpenoid moities through different coupling patterns (Liu et al., 2022). R14, a novel compound isolated from *E. robusta* which has eudesmane-type and diformylated phloroglucinol moieties in our preliminary work was shown an unprecedented EGFR scaffold is presented in this study.

EGFR signaling cascade is a key regulator in cell proliferation, differentiation, division, survival, and disease development. PI3K/AKT/mTOR was a common downstream pathway of EGFR. Renal injury will cause hemodynamic changes, leading to the abnormal activation of PI3K/AKT/mTOR pathway. Besides, TGF-β is thought to be the core factor that contributes to the renal fibrosis. Recently, experimental evidence showed that TGF-β participates in the activation of the PI3K/AKT pathway, suggesting that AKT is a downstream mediator of TGF-β (Li et al., 2015). Thus, TGF-β and AKT may positively crosstalk to regulate each other, promoting tissue fibrosis and other phenotypes. This coincides with our previous work which confirmed that R14 podocyte damage induced by TGF-β (Chen et al., 2022). One research showed *in vitro* experiments in which neutrophils from patients with LN were cultured with curcumin effectively inhibited neutrophil activation under lupus-like conditions, a mechanism potentially mediated via modulation of the PI3K/AKT/NF-κB signalling pathway (Yang et al., 2024). Another study showed that baicalin protected podocytes by downregulating the activity of the PI3K/AKT/mTOR signaling pathway via significantly inhibited the expression of p-PI3K/PI3K, p-AKT/AKT, p-mTOR/mTOR in both *in vivo* and *in vitro* experiments (Ou et al., 2021). Regulation of the above-mentioned pathways is consistent with our results.

AKT, a key downstream effector of PI3K, plays a key role in podocyte apoptosis. Dephosphorylated AKT can increase Bax protein expression and decrease Bcl-2 expression. Bax is a proapoptotic protein and Bcl-2 is an inhibitor of apoptosis protein. The imbalance of Bax and Bcl-2 expressions inducing podocyte apoptosis (Cui et al., 2019). An abundance of evidence indicates that nephrin is important in podocytes both for the slit membrane structure of interpodocytes and the integrity of the filtration barrier (Ettou et al., 2020). Previous data suggested a protective effect of quercetin against podocyte injury through recovering podocytes foot processes with scarce focal fusion and increasing the expression of podocyte markers podocin in the lupus nephritis mice (Dos Santos et al., 2018). Therefore, this experiment aims to prevent and treat LN by preventing podocyte apoptosis, and demonstrates that R14 can improve podocyte injury through marking with podocin and nephrin.

## 5. Conclusion

R14, a novel formyl-phloroglucinol-terpene meroterpenoid isolated from *E. robusta*, features an unusual ring structure derived from the dehydrogenation of isoamyl and a 1’-hydroxyl group. Our previous study established that R14 protected podocytes from undergoing apoptosis. Utilizing the innovative AI model Llama Gram, a new target, the EGFR that is expressed in podocytes was predicted. Subsequent experiments verified this prediction. This is the first time that R14 was discovered to protect podocytes by modulating the EGFR/PI3K/AKT/mTOR signaling pathway. As one of the PTHs, R14 is a novel EGFR scaffold that has never been reported and illustrated to possess this biological property of targeting EGFR. This study clearly lays the theoretical groundwork for further cellular and animal investigations toward ultimate clinical therapies for LN.

## Acknowledgements

Not applicable.

## Materials availability

All of the materials support the conclusions relevant to this manuscript are available upon reasonable request from the lead contact without restriction.

## Declarations

### Ethics approval and consent to participate

Not applicable.

### Consent for publication

Not applicable.

### Authors’ contributions

TC designed the study, performed the data preprocessing and statistical analysis, drafted the manuscript and produced the figures. XL was responsible for flow cytometry experiments. JZ (Juan Zhu) was responsible for cell culture. XZ was responsible for the chemical characterization of R14. JZ (Jianhui Zhang) was responsible for transcriptome sequencing and CMap, HY was responsible for coordinating instruments and funding, XZ was responsible for WB experiments. DW was responsible for molecular dynamics. YZ was responsible for ICC experiments. FZ was responsible for AI models, molecular docking, and bioinformatics analysis. JL conceptualized the study.

